# A scored human protein-protein interaction network to catalyze genomic interpretation

**DOI:** 10.1101/064535

**Authors:** T Li, R Wernersson, RB Hansen, H Horn, JM Mercer, G Slodkowicz, CT Workman, O Rigina, K Rapacki, HH Stærfeldt, S Brunak, TS Jensen, K Lage

**Author notes:** These authors contributed equally (T Li, R Wernersson, RB Hansen). Correspondence should be addressed to Thomas Skøt Jensen and Kasper Lage.

## Abstract

Human protein-protein interaction networks are critical to understanding cell biology and interpreting genetic and genomic data, but are challenging to produce in individual large-scale experiments. We describe a general computational framework that through data integration and quality control provides a scored human protein-protein interaction network (InWeb_IM). Juxtaposed with five comparable resources, InWeb_IM has 2.8 times more interactions (~585K) and a superior functional signal showing that the added interactions reflect real cellular biology. InWeb_IM is a versatile resource for accurate and cost-efficient functional interpretation of massive genomic datasets illustrated by annotating candidate genes from >4,700 cancer genomes and genes involved in neuropsychiatric diseases.

---

Since the turn of the millennium it has become increasingly feasible to experimentally map large-scale protein-protein interaction networks (i.e., hundreds of proteins systematically tested for thousands of interactions in a single study^1-2^). Despite the unquestionable importance of these efforts in humans^3-9^ the most recent screens have only produced in the order of ~25,000 new direct interactions^7,9^, representing only 4-22% of the most conservative estimates of the human interactome^10-11^.

Integration of protein-protein interaction data between heterogeneous databases, different organisms, and from fundamentally different types of interaction experiments is not straightforward. Nonetheless it has been consistently demonstrated that robust computational integration of many different datasets not only improves coverage, but can lead to very high accuracy when the resulting inferred protein networks are tested experimentally^12-13^ or against repositories of well-established interactions (reviewed in 8 and exemplified in 14-16). This is in part because the different experimental large-scale methods complement each other so that no single approach captures the full spectrum of stable and transient interactions between proteins relevant to cell biology^17^. Importantly, the high value of integrated protein networks for the interpretation of vast genomic datasets is illustrated by the onslaught of exome-sequencing projects and genome-wide association studies that have used integrated protein-protein interaction data to reveal non obvious molecular pathways perturbed by somatic mutations in cancers as well as germline common and rare genomic variation in metabolic, psychiatric and immune-mediated diseases (reviewed in 8 and exemplified in 8, 18-22).

We devised a general computational framework to exploit, leverage, and complement ongoing experimental interaction studies and to provide a systematically integrated human protein-protein interaction network for the annotation of genomic and genetic datasets (details can be see in **Methods**, **Supplementary Figure 1** and **Supplementary Table 1**). Specifically, we extracted data from eight heterogeneous protein-protein interaction resources (using only data on physical interactions between proteins or data from protein complexes) covering data from 68,160 independent publications (where independent means articles indexed with a unique PubMed identification number). These data span 4,910,949 redundant protein-protein interactions, 191,336 protein identifiers (covering all accession mapping systems used by the different databases), and stem from 1,493 organisms. Orthology transfer of interaction data is not straightforward and for this reason we used eight different orthology databases with stringent settings and only transferred the interactions if at least four of these databases agreed on the orthology relationship (see **Supplementary Note 1** for an analysis supporting this design). After thorough quality control of the raw data (including filtering to be sure only experimentally measured protein-protein interactions were included in workflow) we created an integrated human protein-protein interaction network named InWeb_InBioMap (InWeb_IM, hereafter).

InWeb_IM consists of 585,843 interactions (**Figure 1 a-h** and **Supplementary Table 1**). When mapping all interactions to UniProt identifiers (IDs) the interactions span 17,530 proteins (or ~87 % of reviewed human UniProt ids). Importantly, 57% of the data come directly from experiments with human proteins, 68% of the data comes from either mice or humans, and 95% of the data comes from human, mouse, rat, cow, nematode, fly, or yeast (**Figure 1e**). We compared InWeb_IM to five widely used human protein interaction networks^23-28^ using a number of quantitative and qualitative metrics (**Figure 1 a-h**, details on other networks are provided in **Methods** and **Supplementary Note 2**). In terms of absolute amount of interactions InWeb_IM has twice the amount of protein-protein interactions compared to I2D, the next-largest network, and 2.8 times the median of all networks (**Figure 1a**). We draw our data from 34.1% more publications than PINA, the network with the next-best coverage in terms of source articles (**Figure 1c**). Considering the total number of proteins that are implicated in one or more interactions, InWeb_IM has 4% fewer than I2D, but more than the remaining four networks (**Figure 1d**). Overall, InWeb_IM has several fold more interaction data than the other networks while also having the largest amount of unique interactions i.e., interactions that are not available from any of the other five networks (344,146 **Figure 1f**).

**Figure 1.**
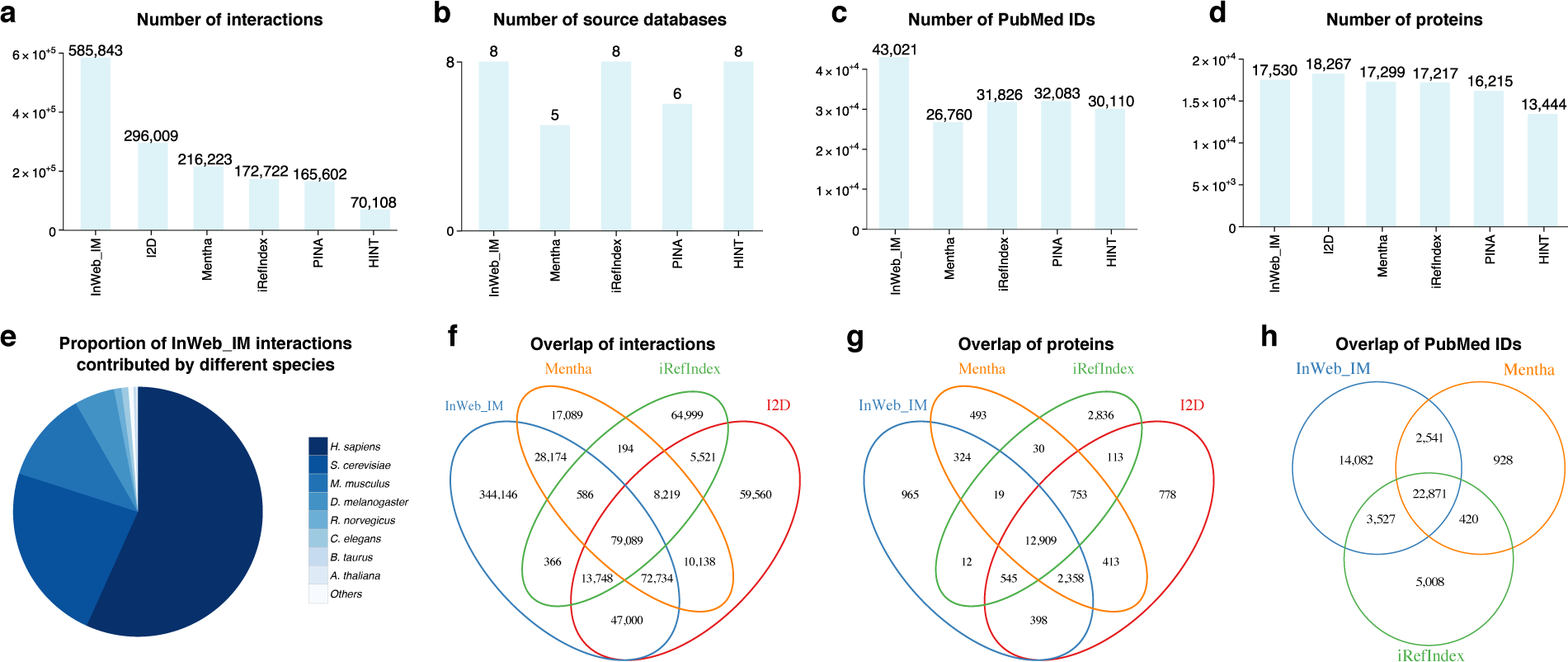
A quantitative comparison of InWeb_IM and five widely used human protein-protein interaction networks. We compared InWeb_IM to Mentha, the protein interaction network analysis platform (PINA), Interologous Interaction Database (I2D), High Quality Interactomes (HINT), and iRefIndex using a number of metrics such as **a)** the total amount of interactions, **b)** number of source databases, **c)** number of PubMed identification numbers, and **d)** number of proteins (using the UniProt reference proteome) covered by at least one interaction. Panel **e)** plots the proportion of InWeb_IM interactions have been found in humans illustrating that 57% (>334,000) of the interactions are not orthology transferred and that 68% (or >398,000) of the interactions have been found in either humans or mice. In the four networks with the most data, we also measured the overlap of **f)** interactions, **g)** proteins covered by at least one interaction, and **h)** publications supporting the interaction data (I2D excluded in panel **c)** and **h)** because PubMed identification numbers could not be traced).

In InWeb_IM interactions are given an initial score based on a number of metrics most notably the reproducibility of the interaction data between different publications (**Methods, Supplementary Figure 1** and **Supplementary Note 3**). To validate the initial score, we defined a highly trusted (gold standard, hereafter) set of protein-protein interactions from pathway databases (**Methods** and **Supplementary Note 3**). We ranked the non-pathway-database-derived interactions based on the initial score and plotted a curve of the enrichment of the gold standard interactions as a function of the rank based on the initial score. This analysis suggests that the initial score indeed up prioritizes the gold standard interactions, as the curve is well above the diagonal and its slope is steeper than the diagonal for the non-pathway-database-derived interactions that rank in the top 30% (**Figure 2a**, **Methods** and **Supplementary Note 3**). We then calibrated the initial score against the gold standard interactions to transform it into a lower bound of the true positive rate of interactions with that initial score or better to give it a probabilistic interpretation (**Figure 2a, Methods** and **Supplementary Note 3**). We confirmed that this final confidence score correlates with an experimentally determined measure of the confidence of interactions between proteins in an independent experiment of 58 immunoprecipitations (**Figure 2c**, **Methods** and **Supplementary Note 4**) with a statistically robust correlation (of 0.38, C.I. [0.35, 0.42]). We repeated this analysis for the only other two network that assigns scores to the interactions, and found a comparable correlation in iRefIndex (0.41, C.I. [0.38, 0.44]), and a lower correlation in Mentha (0.23, C.I. [0.17, 0.29]), where both correlations are statistically significant. Together these analyses confirm the reliability of our score and show it is significantly correlated with an experimentally derived measure of the interaction confidence between proteins.

**Figure 2.**
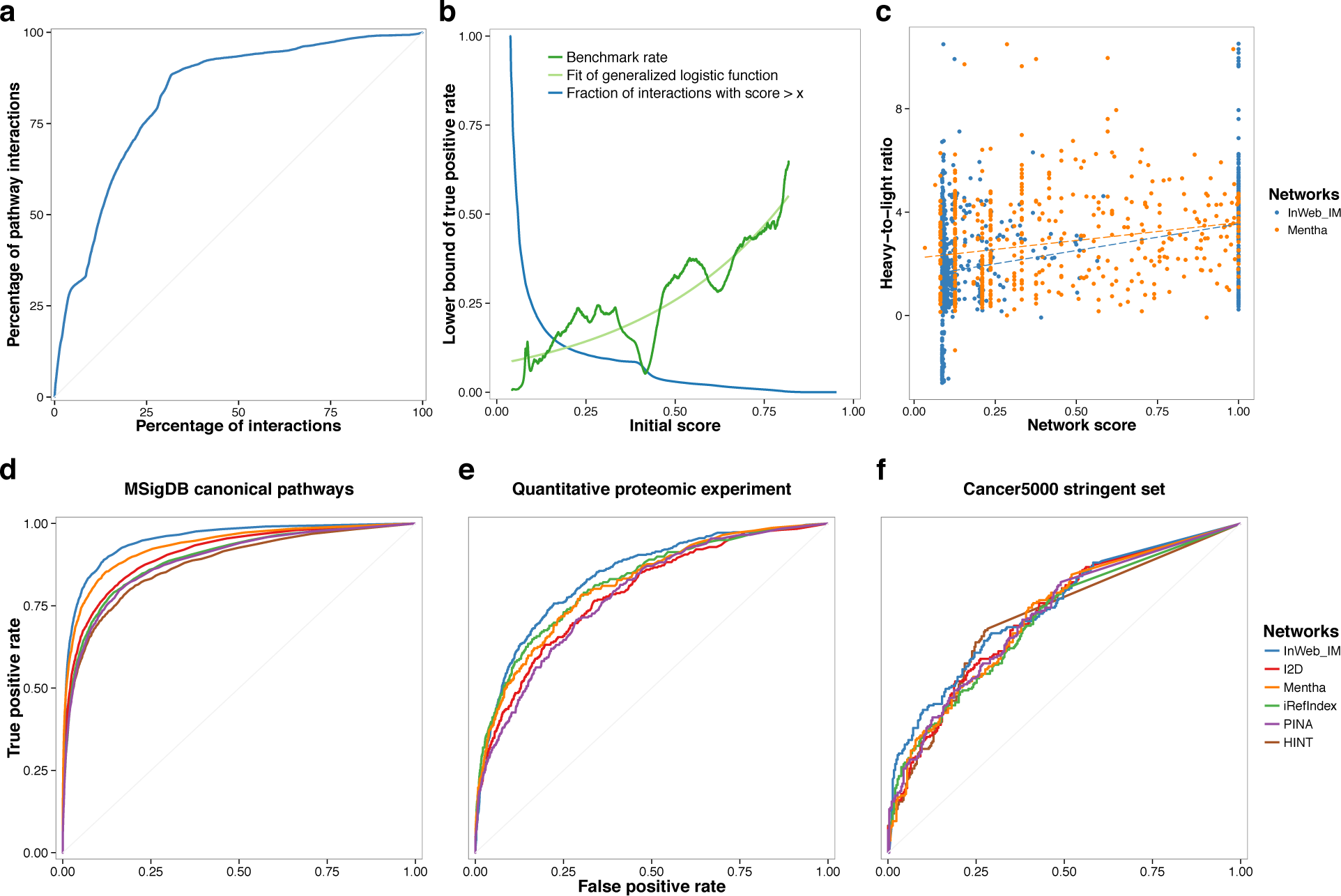
Validating the InWeb_IM score and comparing its biological signal to five other networks. **a**) A plot of the cumulative fraction of the gold standard set (pathway-database-derived interactions) as a function of the rank based on the initial score (normalized to percentages) shows that the initial score prioritizes gold standard interactions (AUC is 0.83). **b**) We calibrated the initial score against the gold standard interactions to show that the initial score correlates with the percent overlap in this set (dark green line for initial data points and light green line for the fitted general logistic function, and the blue line shows the fraction of interactions with scores higher than the values indicated on the x-axis.). Hereby the initial score is transformed into the final confidence score that can be interpreted as a lower bound on the true positive rate (and ranges between 0 and 1). **c)** We observe a significant correlation between the confidence scores from InWeb_IM, Mentha and iRefIndex and experimental values of the confidence of binding between proteins (i.e., the heavy-to-light isotope ratios from mass-spectrometry data of 58 independent human immunoprecipitations). The density of the scores is not uniform across all probabilities because there are fewer interactions with high scores than lower scores for some networks. **d)** We used a cross-validation scheme to test the ability of the six networks to recapitulate pathway relationships between genes in 853 MSigDB canonical pathways through a 30% holdout analysis. Compared to other protein-protein interaction networks InWeb_IM has a superior signal (AUC = 0.95) followed by Mentha (AUC = 0.93); I2D (AUC = 0.91); iRefIndex and PINA (AUC = 0.89), and HINT (AUC = 0.88). **e)** We tested the overall agreement between interactions reported in each network with the aforementioned quantitative protein-protein interactions from 58 immunoprecipitations in human cells. In this analysis, InWeb_IM has the highest AUC (0.84), followed by iRefIndex (0.82), Mentha (0.81), I2D (0.79), and PINA (0.78). **f)** We also tested the ability of InWeb_IM to classify 219 cancer genes from the Cancer5000 Stringent Set defined by Lawrence et al^29^. InWeb_IM has the highest AUC (0.74) followed by I2D, Mentha, and PINA (all 0.72); and HINT and iRefIndex (both 0.71).

The fact that InWeb_IM has many more interactions is not per definition a sign of quality as it could in theory reflect that it is noisier than the other resources. To test the biological signal of InWeb_IM we implemented a classifier and cross-validation scheme (**Figure 2d**, **Methods** and **Supplementary Note 5**) that tests the ability of each network to recapitulate known pathway relationships from 853 stringently defined canonical pathways in the Molecular Signatures Database (MSigDB). We normalized for the amount of proteins covered by data to enable the networks to be compared interaction-for-interaction meaning that we only looked at the signal of the interactions that did exist in each network and did not penalize networks that were smaller than InWeb_IM for missing data (**Methods** and **Supplementary Note 5**). In a 30% holdout analysis InWeb_IM has an area under the receiver operating characteristics curve (AUC) of 0.95 compared to the other networks that range from AUCs of 0.93 to 0.88 with a median of 0.89 (**Figure 2d**). If we do not normalize for coverage, but make an absolute comparison of the ability to recapitulate pathway relationships in MSigDB, InWeb_IM has an AUC of 0.86 which is 16% better than the next-best network (the other five networks range in AUCs from 0.78 to 0.63 see **Supplementary Note 5**) as expected of a high quality network with more than the two times the amount of data than other networks. To further dissect the InWeb_IM data, and to support the quality of both the unique and orthology transferred subsets of interactions in the network, we repeated the analysis on both of these subsets (**Supplementary Note 6**) which resulted in high AUCs (0.90 and 0.85, respectively). These analyses confirm that not only the network as a whole, but also the >344,000 unique interactions and the >252K interactions stemming from orthology transfer have a very good biological signal.

In addition to testing the correlation between the confidence scores from InWeb_IM, Mentha and iRefIndex and the experimentally derived confidence scores from the aforementioned 58 pull downs, we also tested the overall concordance between data in the different networks and this independent set of human protein-protein interactions (comprising 15,205 interactions, **Figure 2e**, **Methods** and **Supplementary Note 4**). Again, InWeb_IM shows the best agreement with this independent dataset (AUC of 0.84) compared to the other networks (AUCs ranging from 0.82 to 0.78 with a median of 0.80).

Many of the genes emerging from recent cancer sequencing studies do not integrate into well-defined pathways and it challenging to functionally interpret the many tumor genomes that are now available. To illustrate the potential for interpretation of massive genomic data sets using InWeb_IM we combined the protein networks with sequencing data from >4,700 tumor genomes (from 21 tumor types) that identified 219 significantly mutated cancer genes^29^ using an algorithm called network mutation burden (NMB). With NMB we tested the ability to predict these 219 cancer genes in a leave-one-out cross validation where genes are classified based on the mutation burden in their first order protein-protein interaction network excluding any information about the gene itself (**Methods** and **Supplementary Note 7**). In essence, this analysis is a fast, data-driven, and systematic way to assign the 219 known cancer genes to networks (i.e., draft pathways) that are associated to cancer based on the overall burden of mutations seen in the network in question. Importantly, this analysis provides contextual information about the molecular groupings of cancer genes. When compared to the five other networks InWeb_IM has the best biological signal (i.e., ability to predict cancer genes, AUC = 0.74 compared to AUCs ranging from 0.72 to 0.71 with a median AUC of 0.72, **Figure 2f**).

To further explore the biological possibilities of the InWeb_IM data we integrated it with tissue-specific expression quantitative trait loci information from the Genotype Tissue Expression Project (GTEx)^30^ to derive 27 protein-protein interaction networks where the corresponding genes are under tissue-specific regulation. On average the tissue-specific networks derived from InWeb_IM have 2-3 times more data than analogous networks derived from the other five resources (**Supplementary Note 8** and **Supplementary Figures 2,3** and **4**). One example is brain, where InWeb_IM has 2.2 times more interaction data connecting brain-regulated genes at the protein level than the next largest network (I2D, **Supplementary Figures 2** and **4**). This observation suggests that it is a particular use of InWeb_IM to discover new pathway relationships in neuropsychiatric diseases.

We tested the ability of each network to annotate and interpret 65 autism genes from a recent study^31^ using NMB, and indeed observe that InWeb_IM is the only network that can assign these autism genes into statistically significant networks with each other (**Supplementary Note 9** and **Supplementary Figure 5**). Furthermore, the quality and applicability of the data for genomic interpretation of diseases with tissue-specific manifestations has further been indicated by the successful use of interim versions of InWeb_IM to interpret genetic data from neurological^19^, cardiovascular^13,32^, immunological^18,33^, metabolic diseases^34^, and cancers^22^. Previous evolutions of the data have also been used as part of the 1000 Genomes Project to annotate population-scale genetic variation^35^.

The curation practices of most protein-protein interaction databases we use are robust, and error rates have been proven to be low (i.e., in the order of ~6%^36^). In addition, the combination of a computationally derived global confidence score and complete transparency of the source and methodology related to each of the 585K interactions provides a multidimensional approach to consolidating the biological relevance of networks and hypotheses derived from InWeb_IM analyses with minimal effort and without the need for particular expertise in protein-protein interaction data or functional genomics networks. This is particularly important for biological follow up of the network analyses because in many cases scrutinizing the original publication(s) and reading the database entry(ies) can also inspire targeted and cost-efficient experiments that ultimately provide proof of specific discoveries and new insight into the cell biology of human diseases (discussed in Ref. 8 and exemplified in Ref. 13).

In addition, the score assigned to each interaction has a probabilistic interpretation, which enables the integration with orthogonal probabilities resulting from genomic and genetic sequencing and genotyping results in Bayesian frameworks. Importantly, the source data for all interactions in InWeb_IM can be easily traced back both in terms of specific publication, database, organism, and experimental method (See **Supplementary Note 10** for an example). This combination of features is unique to InWeb_IM (see **Supplementary Note 11** for a discussion). To exemplify the utility of being able to access the source publications easily and to confirm the high quality of the data in InWeb_IM, we randomly extracted 20 human interactions and read the articles from which the data came (**Supplementary Note 12**). All twenty of the interactions were true positives and for 19 of 20 interactions the databases also had annotated the proteins with correct organism identifiers. In one of 20 interactions the experiments were executed with the murine orthologs of the human proteins and annotated as human in the database, so the interaction was true in the mouse and not a false positive, but an error in the annotation (in terms of organism) of the protein identifiers. This analysis shows that 20 of 20 interactions (100%) were true and 19 of 20 (95%) were annotated with the correct organism identifiers supporting the high quality of the InWeb_IM data and the source database annotations.

InWeb_IM builds on the invaluable foundation laid by the experimental proteomics communities that provide the raw data and the many truly excellent protein-protein interaction databases that heroically extract non-structured data on the physical interactions of proteins from the >25 million articles in PubMed with very high accuracy^36^. We used eight different databases, but the framework presented here can be applied to any number of sources of interaction data. Importantly, interaction screens of particular importance that are not included in any of these resources can be added into the pipeline seamlessly.

While other excellent functional genomics networks like STRING^37^, GeneMANIA^38^ and HumanNet^39^ exist our work focuses on experimental protein-protein interaction data alone (see **Supplementary Note 13** for a discussion of the benefits and drawbacks of protein-protein interaction networks versus other functional genomics networks). Compared to other protein-protein interaction resources, we show that the strengths of InWeb_IM resource lies in a combination of quantitative (e.g., more than double the amount of interactions compared to the next-largest network, >16% better biological signal across 853 pathways, better ability to stratify cancer and autism genes into significant subnetworks and better concordance with an independent human protein-protein interaction dataset) and qualitative features (e.g., a global confidence score with a probabilistic interpretation and transparency in terms of specific publication an interaction originates from). Although some of the other networks also have a good biological signal InWeb_IM is consistently ranked as number 1 (**Figure 2e-f**), while the ranks of the other networks fluctuate. These features make it a versatile resource to interpret and augment very large genomic datasets that are now being produced as part of the ongoing genomic revolution. Our analyses suggest that a few of its uses are to uniquely enable deriving tissue-specific networks e.g., involved in neuropsychiatric diseases and the interpretation of cancer genomes (for example, but not limited to those involved in head and neck squamous cell carcinoma, lung adenocarcinoma, colorectal cancer, and acute myeloid leukemia, **Supplementary Table 2**).

We expect that the general approach we present here to aggregate and score protein-protein interaction data (**Supplementary Figure 1**) - as well as the InWeb_IM network itself - will become increasingly useful with more interaction data and genetic datasets in the future. InWeb_IM is available from https://www.intomics.com/inbiomap. Moreover, we make the data accessible from a graphical user interface http://apps.broadinstitute.org/genets#InWeb_InBiomap so that it can interactively explored by any individual researcher that wishes to study the interactions of proteins of interest. We also provide a roadmap for future updates and overview of file formats and ways to query the InWeb_IM data in **Supplementary Note 14**.

## Online Methods

### Raw interaction data

Raw and partially overlapping interaction data sets were obtained from eight source databases: BIND^1^, BioGRID^2^, DIP^3^, IntAct^4^, MatrixDB^5^, NetPath^6^, Reactome^7^, WikiPathways^8^, along with key information about the individual interaction, such as protein IDs, species, interaction type and PubMed identification numbers for the paper reporting the interaction (where we consider publications independent if they are indexed by a different PMID). Interactions that were annotated as genetic interactions, co localizations, or neighboring reactions were ignored and from the pathway databases (NetPath, Reactome and WikiPathways) we exclusively extracted the small subset of data that describes direct protein-protein interactions or data on experimentally resolved protein complexes. Proteins IDs for all eight databases were mapped to accession identifiers from UniProt^9^, and PubMed identification numbers were used to identify experiments. The authors would like to acknowledge first, that InWeb_IM builds on the invaluable foundation laid by the experimental proteomics communities that provide the raw data and, second, the many high quality protein-protein interaction databases that laboriously extract non-structured data on the physical interactions of proteins from the 25+ million articles in PubMed with a very low error rate^10^.

### Orthology transfer of raw data

Orthology mapping is far from trivial, as different methods rely on different orthology definitions, sequence homology thresholds and handling of paralogs and orthologs. Orthology transfer of interactions in InWeb_IM was built on a voting scheme across eight orthology resources eggNOG^11^, Ensembl^12^, HomoloGene^13^, Inparanoid^14^, Gene^15^, OrthoDB^16^, KEGG^17^, and HOGENOM^18^, where interactions are orthology transferred if four or more databases agree in the orthology assignment (which was determined to give the best biological signal of the resulting network, see **Supplementary Note 1**). Among these inferred interactions we only kept those that were between proteins from the reviewed part of UniProt.

### Calculating confidence scores for the interactions

For each inferred interaction we kept track of the number of publications corresponding to the underlying evidence. Each of these publication contributed to the confidence score for the inferred interaction based on the total number of interactions from the publication; publications describing few interactions contributed more than publications describing many interactions because small-scale experiment are more reliable than interactions from a large screening^19^. In addition, the confidence score was adjusted based on the local topology of the network around the interaction, punishing interactions between proteins with many non-shared neighbors. Finally, using a gold standard set of known high-confidence pathway interactions, the confidence scores were re-calibrated so that a score for an interaction can be interpreted as a lower bound on the probability for the interaction being a true positive. More details and a discussion on how our score fits with the concepts of standardized scoring schemes such as PSISCORE can be found in **Supplementary Note 3**.

### Qualitative and quantitative comparison to other resources

We compared the number of interactions, source databases, supporting publications, and proteins within the above mentioned databases to InWeb_IM. All proteins are indexed using UniProt accessions, which are extracted directly from all networks, and we mapped UniProt to gene symbols using HGNC-provided conversion table for functional analyses. Details can be found in **Supplementary Note 2**.

### Correlating the InWeb_IM confidence score to quantitatively measure protein-protein interactions from an independent experiment

We made a linear correlation of the InWeb_IM confidence scores to experimentally derived quantitative interaction confidences (measured as the heavy-to-light isotope ratios from the mass-spectrometry data of 58 independent human immunoprecipitations using the stable isotope labeling in cell culture [SILAC] method). More details about this metric, the design choice of our experiment, and the 58 immunoprecipitations can be found in **Supplementary Note 4**.

### Quantifying the ability of InWeb_IM to recapitulate pathway relationships

We used an algorithm called Quack (www.broadinstitute.org/genets) to test how well each network can learn structures for 853 stringently defined pathways catalogued in MSigDB normalized for the amount of interactions covered by data in the network being tested. When predicting genes in e.g., the WNT pathway in the 30% holdout analysis, the positive data points were proteins assigned to the WNT pathway MSigDB and the negatives were sampled from the rest of the network. This means that if ten proteins in a pathway (e.g., the WNT pathway) were covered by data in InWeb_IM, but only five of the WNT pathway proteins were covered by data in another network a true positive rate of 100% for InWeb_IM would mean identifying ten out of ten WNT proteins in InWeb, but a true positive rate of 100% for the other network would mean identifying five out of five of the WNT proteins. In this way we are able to determine the biological signal interaction-for-interaction in each network. If we do not normalize for network size, but make an absolute comparison of the ability to recapitulate pathway relationships in MSigDB InWeb_IM has an AUC of 0.86 which is 16% better than the next best network (the other five networks range in AUCs from 0.78 to 0.63). Details can be found in **Supplementary Note 5**.

### Genomic annotation of cancer genes from 21 tumor types

We tested how well known cancer genes can be classified as cancer driver candidates by inferring significance through the aggregated mutation burden of first-order interactors at the protein level using an algorithm Network Mutation Burden (NMB which is described in detail in http://biorxiv.org/content/early/2015/08/25/025445). We used the Cancer5000 stringent set of genes (n = 219) defined by Lawrence et al^20^ as the true positive set and 293 genes that had a FDR = 1 across all 21 tumor types in this paper as negatives. As a negative control for cryptic confounders we randomly selected 87 genes and reran the analysis, which gave a null signal in all networks as expected. More details can be found in **Supplementary Note 7**.

### Concordance between the six networks and an independent dataset of human protein-protein interactions

To evaluate the concordance between an independent set of human protein-protein interactions and the six protein-protein interaction networks, we used both network architectural metrics and quantitative proteomic metrics from the experimental data to test how well predicted physical interactions amongst 58 baits agree with the interactions reported in each of the networks. Specifically, for each resource, we take interactions present in the network as the gold-standard set of known interactions, and constructed a Random Forest model where we used the median-adjusted (by-bait) heavy-to-light ratio, along with Jaccard metric and edge-betweenness centrality, to predict whether each of the 15,205 potential physical interactions are known gold-standard interactions. After training the model on 50% of the data, we computed the Area Under the ROC Curve (AUC) on the remaining 50% of the data (where interactions detected in the experiment are used as positive data points in the analysis and negative data points are interactions not found in the experiment) as a measure of how well interaction data from each network correspond to the quantitative proteomic experiment result. HINT was not included in this analysis because of low overlap between HINT and the experimental data set (less than 1.5% of all of the experimentally derived interactions). More details about the 58 pull down experiments can be found in **Supplementary Note 4**.

### Code and data availability

InWeb_IM and the code for network mutation burden (NMB) will be made available at time of publication from www.lagelab.org. All data will be made public from www.lagelab.org and https://www.intomics.com/inbiomap. Moreover, the data can be accessed from a graphical user interface http://apps.broadinstitute.org/genets#InWeb_InBiomap so that it can interactively explored by any individual researcher that wishes to study the interactions of proteins of interest. We also provide a roadmap for future updates and overview of file formats and ways to query the InWeb_IM data in **Supplementary Note 13**.

## Author Contributions

RW, RBH, TSJ developed the computational framework and continue to maintain InWeb_IM.

TL led and executed the benchmarking analyses and network comparisons with input from RW, RBH, HH, JMM and supervision by KL.

TL, RW, RBH, HH, JMM, GS, CTW, OR, KR, HHS, SB, TSJ, and KL analyzed data.

TL, RW, RBH, TS, KL wrote manuscript with input from all authors.

KL initiated, designed, and led the study.

## Acknowledgements

The authors would like to thank Louis Wich for developing the graphical user interface for InWeb_IM at https://www.intomics.com/inbiomap. TL, HH, JM and KL are supported by a grant from the Stanley Center at the Broad Institute of MIT and Harvard (PI: KL), a Broadnext10 Grant from the Broad Institute of MIT and Harvard (PI: KL), 1R01MH109903 from NIMH (PI: KL) and a grant from the Lundbeck Foundation (PI: KL).

